# Overexpression of α−synuclein in Midbrain Dopamine Neurons Drives Learning Deficits in Mice

**DOI:** 10.64898/2026.11.19.767936

**Authors:** Berenice Coutant, Xiaowen Zhuang, Jillian L. Seiler, Jacob A. Nadel, Kasandra Romo, Yuhong Fu, Robert H. Edwards, Glenda M. Halliday, Talia N. Lerner, Alexandra B. Nelson

## Abstract

Parkinson’s disease (PD) is diagnosed based on core motor features. However, changes in cognitive function are often already present. Cognitive symptoms include changes in “executive function”, including organization, planning, multi-tasking, and flexible decision-making. These early cognitive deficits have been difficult to model and remain understudied. Here, we investigated whether AAV-based overexpression of wild-type human alpha-synuclein (αsyn) in midbrain dopaminergic neurons can produce deficits in learning and behavioral flexibility. We found that αsyn mice develop progressive impairment in basic action-outcome learning. In addition, we found that αsyn mice show impairments in behavioral flexibility during probabilistic reversal learning. Together, these findings demonstrate that αsyn overexpression in midbrain dopaminergic neurons is sufficient to drive cognitive decline. Our early-stage PD model offers a temporal window to dissect cellular and circuits mechanisms underlying non-motor manifestations of PD.

## INTRODUCTION

Parkinson’s disease (PD) is the second most common neurodegenerative disorder. While historically classified by its hallmark motor deficits, cognitive impairment is a common, progressive, and disabling non-motor feature of PD. Cognitive impairment is one of the most important non-motor features of PD and ranges from mild cognitive impairment (MCI) to PD dementia (Aarsland et al., 2021; Carceles-Cordon et al., 2023). MCI can emerge early, at or before clinical diagnosis, and can predict the later development of PD dementia, which occurs in up to 80% of PD patients (Litvan et al., 2012). At the time of clinical diagnosis (when motor signs and symptoms are present) MCI is detectable in 20–35% of patients (Pedersen et al., 2017; Postuma et al., 2012), with deficits most often in executive functions (Dalrymple-Alford et al., 2011; Dirnberger & Jahanshahi, 2013; Mi et al., 2021; Peterson et al., 2009; Tong et al., 2025; Young et al., 2024). Despite its profound functional impact, the onset and underlying neural mechanisms of early cognitive dysfunction in PD remain poorly understood, leaving the field with few effective models to study or therapeutic interventions to prevent or treat cognitive decline.

At the neuropathological level, PD has two hallmarks: abnormal intracellular accumulation and aggregation of alpha-synuclein (αsyn) into Lewy bodies, and progressive dopaminergic neuronal loss in the substantia nigra pars compacta (SNc). Yet the role of αsyn in cognitive impairment remains poorly understood. To date, most preclinical studies exploring the link between αsyn and cognition have relied on transgenic rodent models using pan-neuronal promoters to overexpress wild-type or mutant αsyn globally (Magen & Chesselet, 2011). In these models, cognitive function has been assayed primarily with tests of spatial learning and navigation (Aniszewska et al., 2022; Freichel et al., 2007; Kokhan et al., 2012a; Magen & Chesselet, 2011; Zhou et al., 2008), functions governed by hippocampal and cortical networks, but which rely less on SNc dopamine neurons and their subcortical targets (Chersi & Burgess, 2015). These global overexpression strategies also lack anatomical and cell-type specificity, making it challenging to attribute a cognitive deficit to disease-related processes in distinct brain regions or cell populations, leaving the exact origin of αsyn-mediated cognitive deficits unresolved.

Ambiguity regarding the site of disease pathology that mediates cognitive deficits represents a major translational gap. In clinical populations, early cognitive features include impaired executive functions and manifest as deficits in probabilistic reversal learning and disrupted shifts between goal-directed and habitual control (Cools et al., 2003, 2006; De Wit et al., 2011; Mi et al., 2021; Peterson et al., 2009; Trenado et al., 2018). In people with PD and animal models, these types of cognitive functions are tightly linked to dopaminergic signaling within the nigrostriatal pathway (Cools et al., 2003, 2007; Costa et al., 2015; Klanker et al., 2017; Nuland et al., 2020; Steinberg et al., 2013; Yin et al., 2004, 2005). While all patients presenting with clinical PD exhibit moderate to severe loss of midbrain dopaminergic neurons, αsyn accumulation and dopaminergic cell loss are distinct pathological processes. How and when progressive αsyn pathology *within* dopamine neurons translates into executive dysfunction during the early stages of PD remains unclear.

To address these gaps in knowledge, we investigated action-outcome learning and behavioral flexibility using a cell-type-specific, adeno-associated virus (AAV)-based mouse model overexpressing wild-type human αsyn in midbrain dopaminergic neurons (Barcomb et al., 2025). This model recapitulates the progressive accumulation of Lewy body-like αsyn inclusions and early dopaminergic dysfunction without significant cell death (Barcomb et al., 2025). Crucially, in this model, mice develop only a subtle motor phenotype, which preserves the animals’ capacity to perform complex operant tasks. Therefore, cognitive assays are less likely to be confounded by motor impairment, compared to lesion-based models of PD with severe motor dysfunction. Here, we demonstrate that wild-type human αsyn overexpression in midbrain dopaminergic neurons is sufficient to drive cognitive decline. First, αsyn mice develop progressive impairment in basic instrumental learning. Second, αsyn mice show impairments in probabilistic reversal learning. These findings shed light on the temporal link between αsyn, dopaminergic neurons and the emergence of cognitive dysfunction. Moreover, our model provides a targeted model to dissect the cellular and circuit mechanisms underlying non-motor manifestations of PD.

## RESULTS

### Instrumental learning is progressively impaired following α-synuclein overexpression in midbrain dopaminergic neurons

To assess whether alpha-synuclein (αsyn) overexpression in midbrain dopaminergic neurons induces early cognitive deficits, we overexpressed human wild-type αsyn in dopaminergic neurons of the substantia nigra *pars compacta* (SNc). We randomized DAT-Cre mice to receive bilateral SNc injection of AAVs encoding Cre-dependent αsyn-IRES-mCherry or mCherry alone (Figure 1A-B). Our prior work indicates that this model exhibits early histopathological hallmarks of PD, including reduced striatal tyrosine hydroxylase (TH) expression, increased phosphorylated and aggregated αsyn in midbrain DA neurons, without frank cell loss. In addition, it shows decreased evoked dopamine release and mild locomotor deficits (Barcomb et al., 2025). Consistent with our prior studies, we found that AAV injection produced robust expression of αsyn and mCherry (or mCherry alone) in TH-positive midbrain dopamine neurons (Figure 1B, right panel).

**Figure 1:**
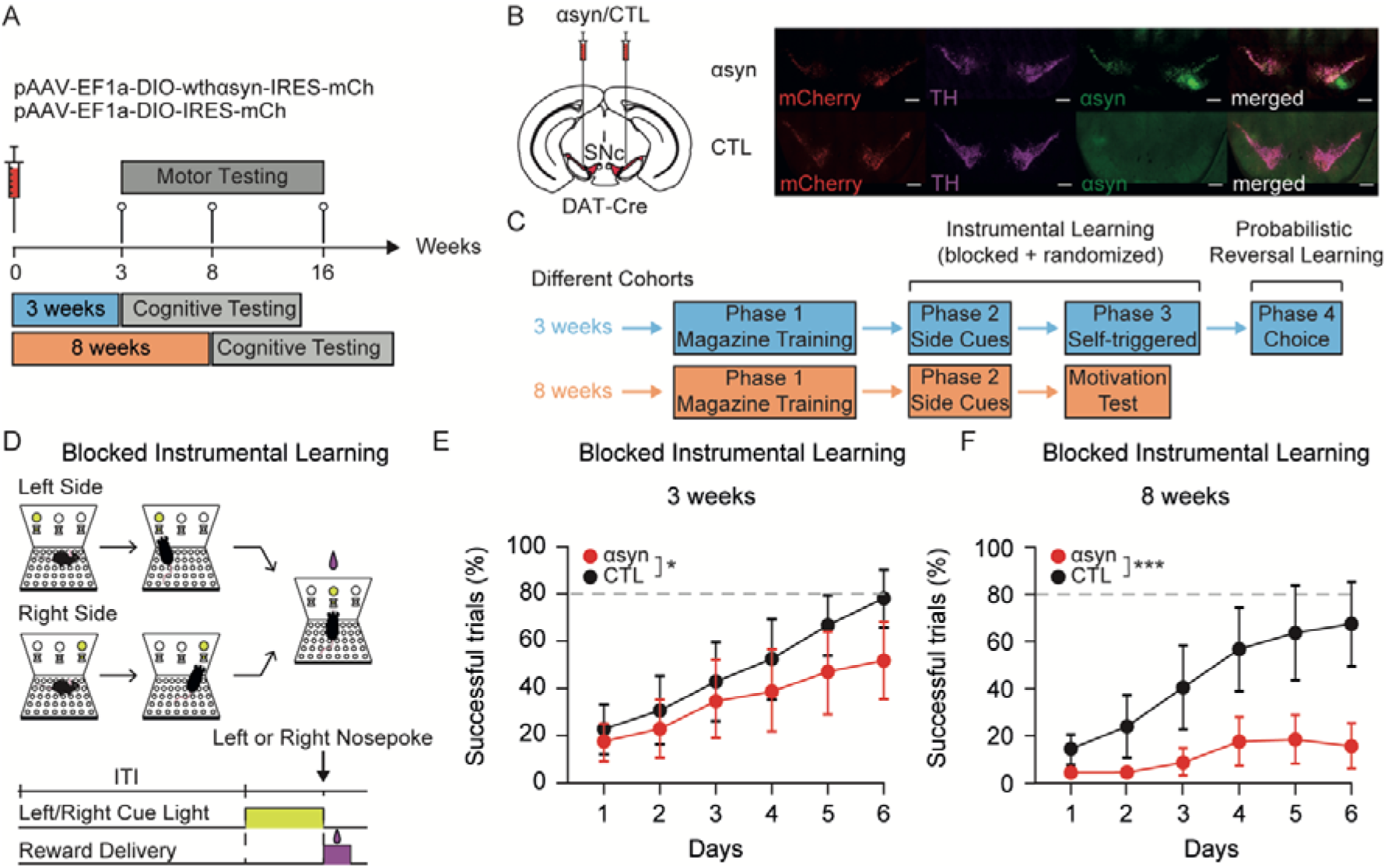
α-synuclein overexpression in midbrain dopamine neurons leads to learning deficits. **A.** Experimental timeline. **B.** *Left:* AAVs encoding human wild-type α-synuclein (αsyn) and/or mCherry (CTL) were injected into the midbrain of DAT-Cre mice. SNc: Substantia nigra *pars compacta*. *Right:* Representative midbrain sections from mice injected with AAVs encoding αsyn and mCherry (top) or mCherry alone (CTL, bottom). From left to right, immunohistochemistry images show mCherry (red), tyrosine hydroxylase (TH, purple), human αsyn (green), and merge. Scale bar: 500 µm. **C.** Cognitive testing timeline. Mice were mildly food restricted during cognitive training and testing periods. **D.** *Top:* Schematic of the Blocked Instrumental Learning Task. *Bottom:* Trial structure. ITI: intertrial interval. In this task, a cue appears above the left or right port. If mice nosepoke at that port, reward is delivered to the central reward port. Cues appear above the left or right port in blocks. **E-F.** Successful trials (%) across 6 training days in αsyn and CTL mice, at 3 weeks (E, n=19, 20) or 8 weeks post-injection (F, n=20, 13). A linear mixed-effects model showed a significant effect over time at 3 weeks (Day: p = < 2e^-16^) and 8 weeks (Condition × Day: p = 9.480×10 ^-09^). There were also significant differences between conditions over time (Condition × Day: p = 0.0338). On Day 6, αsyn mice have fewer successful trials than CTL mice (Day 6: αsyn vs CTL: p = 0.0165). Dotted grey line indicates the inclusion/learning criterion. Data are presented as mean ± SEM. See Supplementary Information, Table 1 for statistics.

To determine the cognitive impact of cell type-specific expression of αsyn in dopaminergic neurons, we used an operant conditioning paradigm (Figure 1C–D) assaying action-outcome, or instrumental, learning. Animals were food-restricted (seMethods for details) and placed in an operant chamber containing a center reward port and two side choice ports (Figure 1D). During each trial, a light cue appeared above and within one side port to indicate the reward-associated side. If the mouse nosepoked the illuminated side port, light cues would extinguish, and food reward would be delivered to the center port. Task sessions were initially organized in blocks of trials rewarding only one side choice (“blocked”), and later blocks contained a mixture of trials with rewarded left or right choices (“randomized”). Behavior and subsequent analysis were performed blinded to the AAV group.

We assessed behavior on the blocked task at 3 or 8 weeks after AAV injections, in separate cohorts of mice. At 3 weeks post-injection, both αsyn-mCherry (αsyn) and mCherry alone (control, CTL) mice showed gradual improvements in performance over 6 days of training (Day: p<0.0001; for all statistical details, see Supplementary Table 1), with a small, but significant difference between groups over time (Condition × Day: p=0.034; Figure 1E, left panel). These findings are consistent with preserved but modestly slowed learning in αsyn mice at this stage. At 8 weeks, mCherry control mice showed a similar pattern of learning over 6 days (Figure 1F). In contrast, αsyn mice failed to acquire even simple action–outcome associations at this timepoint (Condition × Day: p<0.0001; Figure 1F). These results indicate that αsyn overexpression in midbrain dopamine neurons leads to progressive impairment in instrumental learning.

### Cognitive Decline Is Paralleled by an Intact Reaction Time, but a Progressive Lack of Motivation

Operant performance depends on multiple components, including instrumental learning of action-outcome associations (“cognitive”), motivational value of the outcome (“motivation”), and motor execution, which can be dissociated experimentally (Dickinson & Balleine, 1994; Yin et al., 2006). The progressive decline in instrumental learning observed in the AAV-αsyn model could be confounded by deficits in motor function and/or motivation. To determine whether this deficit reflects cognitive dysfunction rather than motor or motivational impairments, we analyzed additional behavioral metrics. Reaction time, measured as the latency between side cue and side nosepoke (when it occurred), was unchanged between groups at both 3 (Condition × Day: p=0.528; Figure 2A, left panel) and 8 weeks (Condition × Day: p=0.587; Figure 2A, right panel), arguing against motor impairment as a driver of the observed deficits. This was further supported by our independent motor assessments (Supplementary Figure 1A–D), which revealed a modest and stable reduction in locomotor velocity in αsyn mice (Supplementary Figure 1A), in contrast to the progressive impairment in instrumental learning (Figure 1E-F). Performance on the rotarod and pole test was comparable between αsyn and control mice over time (Condition × Age: p=0.929; Supplementary Figure 1B; Condition × Age: p=0.779). These findings are consistent with our previous characterization of this model (Barcomb et al., 2025). The number of central nosepokes, reflecting combined motor and motivational drive, was similar at 3 weeks (Condition × Day: p=0.204; Figure 2B, left panel) but reduced in αsyn mice at 8 weeks (Condition: p<0.01; Figure 2B, right panel), suggesting a potential motivational impairment contributing to the profound deficit in learning observed at 8 weeks.

**Figure 2:**
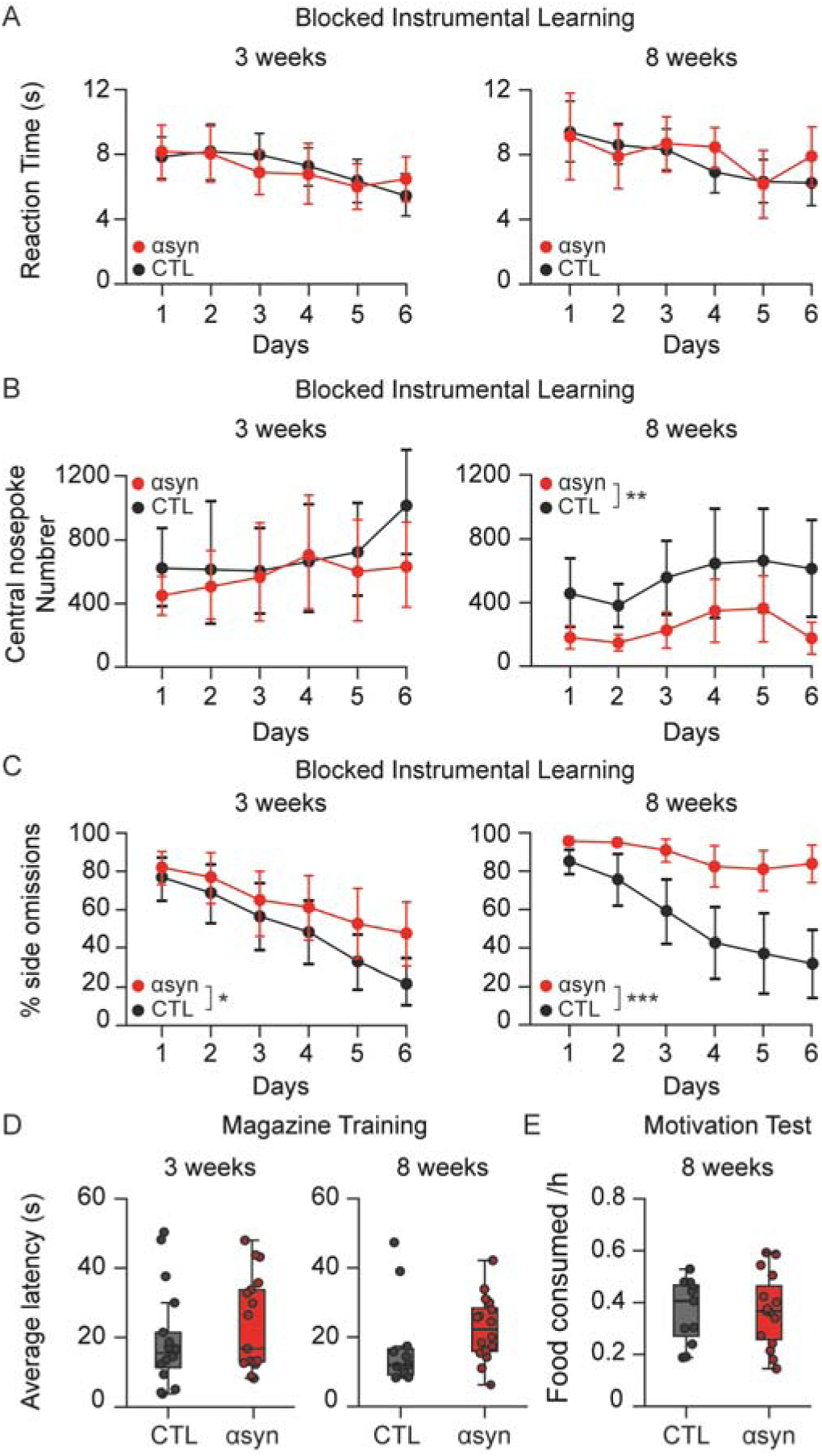
Learning deficits are paralleled by increased omissions. **A.** Reaction times in the blocked Instrumental Learning Task. Reaction time (s) at 3 weeks (left) or 8 weeks post-injection (right) was defined as the time between side cue onset and side nosepoke, when a side nosepoke occurred. Reaction time did not differ between αsyn and CTL mice at 3 weeks (Condition × Day: p = 0.5283, n=19, 20), nor at 8 weeks (Condition × Day: p = 0.5871, n=20, 13). **B.** Task engagement in Blocked Instrumental Learning Task. The number of central nosepokes in αsyn and CTL mice at 3 weeks (left) or 8 weeks (right) after AAV injections. The number of central nosepokes did not differ between groups at 3 weeks post-injection (Condition: p = 0.4771, n=19, 20) but was significantly different at 8 weeks (Condition: p < 0.01, n=20, 19). **C.** Omissions (defined as failure to nosepoke in a side port within 10 seconds of cue onset) in the Blocked Instrumental Learning Task. Side omissions (%) at 3 weeks (left) or 8 weeks (right) post-injection. At 3 weeks, side omissions were increased in αsyn versus CTL mice over training days (Condition × Day: p = 0.0390; n=19, 20), specifically on day 6 (Day 6: αsyn vs CTL: p = 0.0180). At 8 weeks, side omissions were also increased in αsyn mice (Condition × Day: p < 0.0001), specifically on days 2 to 6 (n=20, 13). **D.** Latency to retrieve reward (s) during initial magazine training in αsyn and CTL mice at 3 weeks (left) or 8 weeks (right) after AAV injections. Latency did not differ between αsyn mice at 3 weeks (Welch’s two-sample *t*-test: p = 0.4191; n=20, 22), nor at 8 weeks (Welch’s two-sample *t*-test: p = 0.5598; n=16, 12). **E.** Food consumed per hour in αsyn and CTL mice 8 weeks after AAV injections. Food-restricted mice were given unlimited access to sugar pellets for one hour. The amount of food consumed (g/hour) did not differ between αsyn (n = 15) and CTL (n = 11) mice (Unpaired two-tailed Student’s *t*-test: p = 0.923).

We next examined side omissions (failure to poke a side port during the cued choice period), a proxy for task understanding in this paradigm. If mice do not understand they have to follow side cues and nosepoke at side ports, they may only nosepoke at the center in an effort to obtain reward. At 3 weeks post-injection, both groups showed a decrease in the percentage of side omissions over time (Day: p<0.0001; Figure 2C, left panel; Supplementary Information, Table 1), consistent with successful learning. However, the amplitude of this decrease differed between groups (Condition x Day: p=0.039; Figure 2C, left panel), with αsyn mice showing a marginally higher rate of side omissions. Control mice tested at 8 weeks also showed decreasing side omissions across learning, while αsyn mice showed persistently high numbers of side omissions over training days (Condition x Day: p<0.0001; Figure 2C, right panel), paralleling their failure to acquire the task (Figure 1E, right panel).

Finally, in a cue-independent “magazine training” condition, where food rewards were delivered to the center port periodically, reward retrieval latency was unchanged at 3 or 8 weeks in αsyn compared to mCherry controls (p=0.419 and p=0.560, respectively; Figure 2D), consistent with preserved motivation and reward retrieval between groups across time. In the 8-week post-injection groups, we also tested the overall motivation for food, by comparing the amount of sugar pellets consumed in one hour (g/h) in the home cage. At 8 weeks, mCherry controls and αsyn mice consumed similar amounts of food (p=0.923; Figure 2E). Together, these results indicate that the learning deficits observed in αsyn mice are unlikely to be explained by alterations in food motivation, reward retrieval, or motor performance.

### Overexpression of α-synuclein Impairs Instrumental Learning in a Subset of Mice at 3 Weeks

The severe learning impairment we saw at 8 weeks of αsyn overexpression prevented the study of more complex instrumental learning tasks (“randomized’) and probabilistic reversal learning. At 3 weeks, however, mice that had met criteria on the “blocked” task could be transitioned to the more cognitively demanding “randomized” version of the instrumental learning task. In the randomized instrumental learning paradigm, animals were required to self-initiate trials with a center nosepoke, at which time a side choice cue was provided either on the left or right side, in a randomized fashion (Figure 3A). With a nosepoke on the cued side, reward was delivered to the center port. In the randomized instrumental task, αsyn mice that met inclusion criteria performed similarly to mCherry controls (Condition x Day: p=0.634; Figure 3B, upper panel).

**Figure 3:**
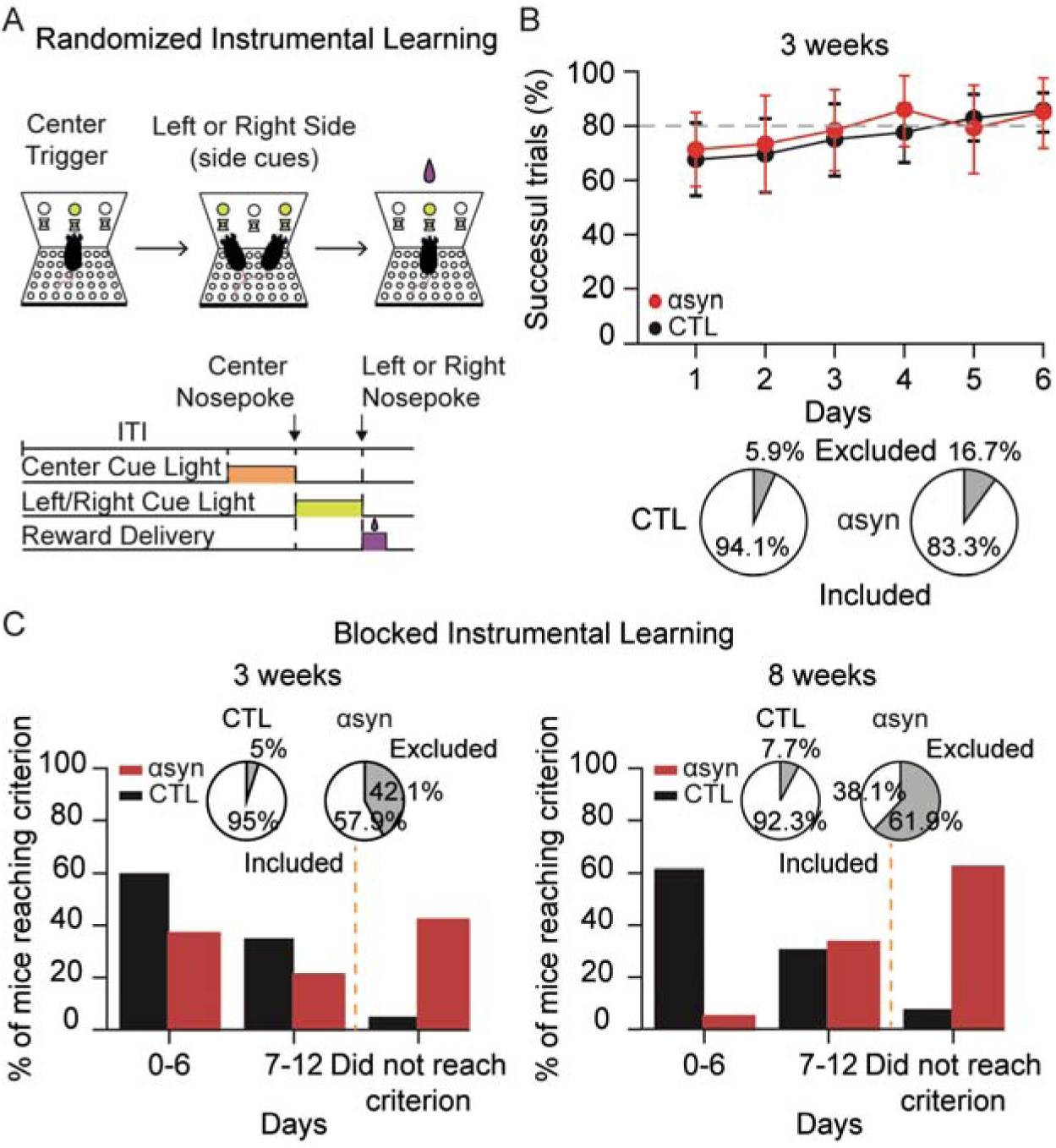
α-synuclein Overexpression leads to subtle learning deficits at 3 weeks post-injection. **A.** Top: Schematic of the Randomized Instrumental Learning Task. Bottom: Randomized Instrumental Learning trial structure. ITI: intertrial interval. **B.** Learning curve showing successful trials (%) in αsyn and CTL mice over 6 days of training, at 3 weeks post-injection. Performance did not differ significantly between αsyn and CTL mice over time (Condition × Day: p = 0.634; n=12,17). Dotted grey line represents learning criterion. Data are presented as mean ± sem. Bottom: Pie charts representing % of excluded (light grey) and included mice in CTL (dark grey, left) and in αsyn (red, right). **C.** Blocked Instrumental Learning Task. Bar plots showing the % of mice reaching criterion (80% successful trials in max. 12 days) over time at 3 weeks (left) and 8 weeks (right) post-injection. Pie charts indicate the fraction of mice that did not meet criterion (excluded; grey) 3 weeks (left) and 8 weeks (right) after AAVs injections. Vertical dotted orange line represents the criterion.

In addition to differences in performance between αsyn and mCherry control mice over the first 6 days of training on the blocked task (Figure 1E), a substantial proportion of αsyn mice failed to reach the learning criterion within 12 days on the blocked task (αsyn: 8/19 (42%) vs CTL: 1/20 (5%) excluded at 3 weeks; αsyn: 13/20 (62%) vs CTL: 1/13 (7%) excluded at 8 weeks; Figure 3C). Consequently, many mice were excluded from subsequent training and testing. These findings suggest that even at early stages, αsyn overexpression impairs action-outcome instrumental learning in a subset of animals, which is masked when only high-performing individuals are analyzed.

### Overexpression of α-synuclein in Midbrain Dopaminergic Neurons Impairs Behavioral Flexibility in Probabilistic Reversal Learning

To overcome this roadblock in studying more complex forms of learning and decision-making, we implemented a simplified training paradigm to promote learning in a new cohort of mice. In this version of early training, cue lights appeared only inside both side ports, nosepokes at either side port led to reward delivery, and no time limit was enforced between the trial start and side port nosepokes (Figure 4A-B). This modification of the task enabled all animals to meet criterion (80% successful trials in <12 days) and proceed to more challenging tasks. Although all mice reached criterion, αsyn mice did require more time to reach criterion than mCherry controls (p=0.028; Figure 4C), indicating that αsyn overexpression slows acquisition.

**Figure 4:**
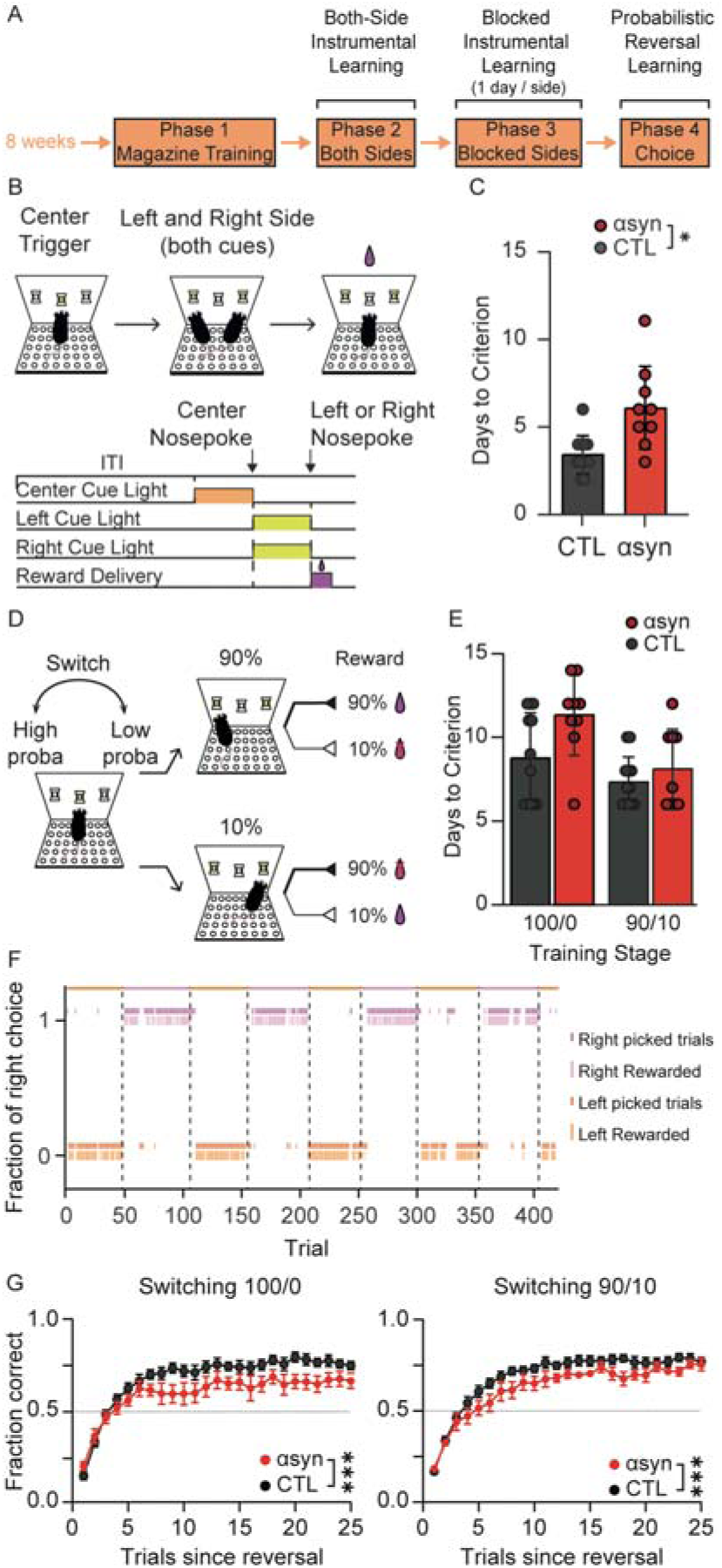
Overexpression of α-synuclein Leads to Deficits in Reversal Learning. **A.** Cognitive testing timeline. **B.** *Top:* Schematic of the Both-Side Instrumental Learning Protocol. *Bottom:* Both-Side Instrumental Learning Task structure. ITI: intertrial interval. In this task, a cue appears above the central port, where mice poke for trial initiation. Then cues appear above both left and right ports. If mice nosepoke at either side port, reward is delivered to the central reward port. **C.** αsyn mice took longer than CTL mice to reach to criterion (80% successful trials) on the Both-Side Instrumental Learning task (Bonferroni–Šídák-adjusted post hoc comparison: p = 0.0278; n=9, 12). **D.** Schematic of Probabilistic Reversal Learning Task (90/10 task is illustrated). **E.** αsyn and CTL mice took a similar number of days taken to reach criterion (>80% accuracy over 3 consecutive days for 100/0 and >70% accuracy for 90/10 contingencies) on the Reversal Learning Task, under both 100/0 (Condition: p = 0.0943 n=9, 12) and 90/10 contingencies (Condition: p = 0.7879; n=9, 12). **F.** Diagram illustrating reversal learning behavior over a representative 90/10 reversal learning session of 400 trials. The colored bars at the top of the diagram indicate the high probability port for that block (orange = left; purple = right), which was switched after 50 trials with a nosepoke in the higher probability port. Dark orange ticks represent left choices (nosepokes), whereas light orange ticks represent rewarded trials associated with left choices; dark purple ticks represent right choices (nosepokes), whereas light purple ticks represent rewarded trials associated with right choices. **G.** Fraction of choices of the high reward probability port after contingency reversal in the 100/0 (left) and 90/10 (right) Probabilistic Reversal Learning Task. Performance was lower in αsyn versus CTL mice on both 100/0 (Trials x Condition: p < 0.0001; n=9, 12) and 90/10 versions of the task (Condition: p = 0.0416; n=9, 12).

We then assessed behavioral flexibility using a probabilistic reversal learning task. In this task, mice self-initiated each trial with a center nosepoke. After trial initiation, both side cues were presented, but one port was associated with a high probability of reward and the other with a low probability. Mice were first trained under a deterministic 100/0% reward probability schedule for a maximum of 14 days, with the contingencies switched every 40 rewarded trials. Later, the task was made more challenging by introducing uncertainty with a 90/10% reward schedule, with contingencies switched every 50 trials with choices on the high reward probability side (Figure 4D). Mice overexpressing αsyn learned reward contingencies under both deterministic (100/0) and probabilistic (90/10) conditions similarly to mCherry controls (100/0: p=0.094; 90/10: p=0.79; Figure 4E). A representative session of reversal learning behavior in a well-trained mouse is shown in Figure 4F. However, following contingency reversal, αsyn mice showed a significant impairment in adapting their choices, with fewer choices of the high reward-probability port compared to mCherry controls under both deterministic and probabilistic conditions (100/0: Trials x Condition: p<0.0001, Figure 4G, left panel; 90/0: Condition: p=0.042, Figure 4G, right panel). These results indicate that αsyn overexpression in midbrain dopaminergic neurons impairs behavioral flexibility as measured in a probabilistic reversal learning task.

## DISCUSSION

To study the role of alpha-synuclein (αsyn) in early cognitive deficits seen in Parkinson’s disease, we used an AAV-based αsyn overexpression mouse model. We found that cell type-specific overexpression of human wild-type αsyn in midbrain dopaminergic neurons was sufficient to drive progressive deficits in action-outcome, instrumental learning and behavioral flexibility. At 3 weeks of αsyn overexpression, mice showed subtle impairments, but by 8 weeks these mice showed a profound impairment in acquisition of a basic instrumental learning task. Moreover, in a reversal learning task, we found αsyn mice were impaired in adapting to a new reward contingency. Altogether, these findings identify progressive alterations in instrumental learning and behavioral flexibility as early cognitive consequences of αsyn dysfunction in midbrain dopaminergic neurons.

Cognitive impairment is a complex and disabling non-motor feature of PD, responding poorly to dopamine replacement therapies (Dirnberger & Jahanshahi, 2013). The only currently approved therapy, cholinesterase inhibitors, provide a modest symptomatic benefit to those with Parkinson’s and dementia (Emre et al., 2004; Mamikonyan et al., 2015; Van Laar et al., 2011), suggesting that additional mechanisms may contribute to cognitive dysfunction. Cognitive impairment is common and can occur at or before clinical diagnosis, meaning before motor deficits emerge (Pedersen et al., 2017; Postuma et al., 2012). People with PD often develop dementia later in disease progression (Janvin et al., 2005, 2006). As those who develop early cognitive symptoms are at high risk for developing dementia, it is vital to understand the mechanisms of cognitive impairment for early intervention.

Cognitive impairment in people with PD includes frontal/executive dysfunction (Dirnberger & Jahanshahi, 2013; Ryman et al., 2021; Tong et al., 2025). Executive function encompasses organization, planning, decision-making, and the ability to be flexible across different contexts. Clinical studies have indeed found that people with PD struggle to adapt to shifting reward contingencies (Ciafone et al., 2020; Peterson et al., 2009; Trenado et al., 2018). Deficits in goal-directed control, one component of decision-making, using an instrumental learning task, have also been reported in people with PD (Mi et al., 2021). Our finding that αsyn overexpression in midbrain dopaminergic neurons is sufficient to impair instrumental learning and behavioral flexibility mirrors observations in people with PD and mild cognitive impairment.

In our AAV model, we found subtle learning impairment at 3 weeks and profound impairment at 8 weeks, suggesting a cognitive decline over time. Operant behavior must be interpreted considering potential confounds, such as motor impairment or loss of motivation. We found very mild motor signs at 3 and 8 weeks: subtle decreases in open field locomotor velocity, but no changes in rotarod or pole test performance (Supplementary Figure 1B-D). This is consistent with our previous report (Barcomb et al., 2025). The progressive learning impairment we observed did not align with progressive motor deficits. Indeed, motor deficits were mild and stable over time on this model. Consistent with overall preserved motor function, here we found that αsyn mice showed normal reaction time in the operant box. We did identify some signs of mildly reduced motivation, which we tested with several assays. Mice overexpressing αsyn showed more side (choice) omissions during instrumental learning, which could reflect cognitive (poor understanding of the task) and/or motivational deficits. However, more direct assessments of general motivation for food rewards (latency to obtain free rewards in the operant box and consumption of sugar pellets in the home cage) did not show significant differences. Together, these findings suggest motor and motivational deficits are mild and unlikely to be the primary drivers of the poor instrumental learning observed.

Interestingly, in our reversal learning paradigm, the pattern of impairment differed according to reward uncertainty. Under deterministic contingencies (100/0), αsyn mice showed a lower asymptotic level of performance, whereas under probabilistic contingencies (90/10), they showed slower initial adaptation but ultimately approached control performance. This pattern differs in some respects from observations in people with PD, who can show impairments in both probabilistic learning and reversal (Peterson et al., 2009; Trenado et al., 2018). Differences in disease stage, extent of dopaminergic degeneration, and task structure may contribute to these differences. Our model provides an early, pre-degenerative window where dopamine release is compromised but structural pathways remain intact.

### Learning impairments relate to dopamine dysfunction and occur before cell death

Prior work studying cognitive function in αsyn mouse models of PD has often focused on spatial learning and memory, using tasks such as the Morris Water Maze (Freichel et al., 2007; Kokhan et al., 2012b; Nuber et al., 2008; Zhou et al., 2008). The Morris Water Maze is widely used to assess hippocampal-dependent spatial learning and memory (Da Cunha et al., 2003). However, spatial learning and memory are less likely to be impacted in early-stage PD. In autopsy-confirmed cases of cognitive impairment with αsyn pathology, without Alzheimer’s disease co-pathology, executive dysfunction was the primary cognitive manifestation (Ryman et al., 2021). Fewer studies in rodent models of PD have focused on tasks relevant to executive dysfunction. In healthy animals, instrumental learning and behavioral flexibility engage nigrostriatal circuits and are strongly modulated by dopaminergic signaling (Balleine et al., 2009; Graybiel, 2008; Howard et al., 2017; Lerner et al., 2015; Morris et al., 2006; Parker et al., 2016). Reversal learning impairment was observed in a pan-αsyn transgenic mouse model (Magen et al., 2012). However, the lack of spatial specificity of αsyn overexpression in this model make it difficult to determine which brain area is responsible for this effect. By establishing a cell-type-specific, AAV-based mouse model, we were able to bypass the severe motor confounds of classic toxin-based models like 6-Hydroxydopamine (6-OHDA) and 1-méthyl-4-phényl-1,2,3,6-tétrahydropyridine (MPTP) and the anatomical ambiguity of global transgenic lines. A recent AAV-based study also reported impaired reversal learning following αsyn overexpression (Decourt et al., 2023), although that study used the A53T mutant form of human αsyn, a pathogenic variant associated with early-onset familial PD (Lashuel et al., 2012). Our use of wild type αsyn provides complementary evidence that increased expression of protein, rather than a highly pathogenic familial mutation, can disrupt these behaviors. Selective αsyn overexpression in midbrain dopaminergic neurons is highly relevant in early PD pathology (Surmeier et al., 2017).

Our previous work in this AAV-based αsyn overexpression model showed that overexpression caused αsyn pathology in midbrain dopamine neurons, very modest motor dysfunction at 2 months, but no overt cell loss at up to 4 months after injection (Barcomb et al., 2025). However, we found a significant reduction in evoked striatal dopamine release. Taken together with the present study, our findings suggest that significant cognitive impairment can develop at or even before motor dysfunction, mirroring observations in people with PD. Given the cell type specificity of our manipulation, this finding also suggests that some aspects of cognitive impairment in PD may be mediated by alterations in dopaminergic cell function, even before overt cell loss or involvement of other brain regions. Pathological αsyn accumulation has been shown to physically disrupt vesicular trafficking and clustering at the active zone (Vargas et al., 2017), impair dopamine transporter distribution and expression (Bellucci et al., 2011), and suppress striatal long-term potentiation (Tozzi et al., 2016) before structural neurodegeneration (Janezic et al., 2013). Our findings align with the idea that phasic, point-to-point dopamine is important for learning while slower tonic signals are adequate to support gross movement (Cai et al., 2024; McGregor et al., 2026).

Our cell-type specific, AAV-based, αsyn mouse model of PD offers a unique pre-degenerative window for investigating the precise molecular and cellular mechanisms responsible for early cognitive dysfunction before overt neurodegeneration. However, there are several caveats and limitations to these findings. First, the severe impairment in basic instrumental learning that we identified at 8 weeks of overexpression presents an important limitation for the testing and interpretation of more complex operant behaviors. Such tasks depend on acquisition of the initial instrumental contingency. We were able to address this limitation by making the initial training procedure simpler and more permissive, which helped animals to learn enough to proceed to the reversal learning task. Nevertheless, this issue highlights the importance of distinguishing impairments in task acquisition from impairments in the specific cognitive process targeted by a more complex task. Future studies might perform basic operant training prior to AAV administration to work around this issue. Second, although in-task measures of motor function (such as reaction-time) did not differ between groups, other measures indicated the potential for reduced motivation (decreased number of central nosepokes at later stages). Third, our experiments also address only a subset of the cognitive domains affected in PD and do not establish whether αsyn overexpression alters other executive processes, episodic memory, or the balance between goal-directed and habitual control. Finally, our model targeted both SNc and ventral tegmental area dopaminergic neurons, preventing attribution of the phenotype to a single midbrain population. However, this broader midbrain targeting may also have translational relevance, given that αsyn pathology can affect multiple dopaminergic nuclei in PD despite differences in their vulnerability to neuronal loss (Aarsland et al., 2021; Surmeier et al., 2017). Future studies using more spatially and anatomically resolved manipulations will therefore be important for determining the relative contributions of distinct dopaminergic populations and other PD-vulnerable regions to early cognitive dysfunction.

Taken together, our findings demonstrate that human wild-type αsyn overexpression in midbrain dopaminergic neurons is sufficient to produce progressive deficits in instrumental learning and behavioral flexibility before severe motor dysfunction and overt neurodegeneration.

The progressive emergence of cognitive dysfunction over time, together with substantial heterogeneity at earlier stages, suggests that αsyn pathology does not trigger an immediate behavioral phenotype, but rather leads to dysfunction gradually. This temporal progression provides an experimental window for investigating why some dopaminergic circuits become functionally vulnerable to αsyn pathology before overt cell death, and how they might contribute to specific behavioral phenotypes. More broadly, this cell-type specific model provides a tool for investigating the cellular and circuit mechanisms that determine vulnerability to early cognitive dysfunction, as well as those cellular or circuit adaptations that help maintain normal function.

## METHODS

### Animals

To target midbrain dopaminergic neurons, three-month-old hemizygous DAT-IRES-Cre mice (B6.SJL-*Slc6a3^tm1.1(cre)Bkmn^*/J or DAT^IRES*cre*^; Strain No. 006660, RRID IMSR_JAX:006660, Jackson Labs) were crossed to wild-type C57Bl/6J (Strain No. 000664, RRID IMSR_JAX:000664, Jackson Labs) to yield hemizygous DAT-Cre mice. Animals were housed 1–5 per cage on a 12 hour light/dark cycle with *ad libitum* access to food and water unless food restricted (see below). When food restricted, animals had limited access to food and *ad libitum* access to water. All experiments were performed during the light phase. We complied with local and national ethical regulations regarding the use of mice in research. All experimental protocols were approved by the University of California San Francisco and/or Northwestern University Institutional Animal Care and Use Committee.

### Surgical procedures

A detailed surgery protocol (DIO: dx.doi.org/10.17504/protocols.io.dm6gpdw1dgzp/v1) can be found at https://www.protocols.io/view/mouse-stereotaxic-surgery-dm6gpdw1dgzp/v1. Briefly, anesthesia was induced with either isoflurane (3%) or intraperitoneal (I.P) injection of ketamine (40 mg/kg) / xylazine (10 mg/kg) and maintained with 0.3%-0.8% inhaled isoflurane. Mice were placed in a stereotaxic frame and a mounted drill was used to create holes over the substantia nigra *pars compacta* (SNc). To model Parkinson’s disease in mice, 300 nL of undiluted AAV encoding wild-type human alpha-synuclein (αsyn) and mCherry, AAV5-Ef1a-DIO-wthasyn-IRES-mCh (Vector Builder, titer 3.83×10^13^ GC/ml, lot #211030AAVJ01) was injected bilaterally in SNc (-3.1 AP; -2.2 ML; -4.1 DV). In control mice (CTL), 300 nL of undiluted AAV encoding mCherry alone, AAV5-Ef1a-DIO-IRES-mCh (Vector Builder, 3.95×10^13^ GC/ml, lot #211104AAVD11), was used. Viruses were injected at 50 nL per minute with a microinjector (Micro4, WPI), and the injection syringe was left in place for 10 minutes prior to being withdrawn, and the scalp sutured.

### Motor assessment

Details of the accelerating rotarod test and open field test can be found dx.doi.org/10.17504/protocols.io.q26g7yo4kgwz/v1 at https://www.protocols.io/private/F4255C85C47B11EEB0C50A58A9FEAC02. Briefly, locomotion and motor coordination were evaluated using the open field, accelerating rotarod tests (Ugo Basile) and pole test (Kim et al., 2019) at three time points: 3, 8, and 16 weeks after αsyn-mCherry (αsyn) or mCherry alone (CTL) AAV injections. Prior to the first testing session, mice were habituated for 2 days, 30 minutes each, in the open field (clear acrylic cylinders, 25 cm in diameter, Tap Plastics). During each experimental session, movement was monitored with an overhead camera and analyzed offline using video-tracking software (Noldus Ethovision), including distance traveled, velocity and rotational behavior (90° contralateral or ipsilateral turns). To minimize the effect of motor learning on rotarod performance, each mouse underwent only one session per timepoint. One session included three trials with a 5-minute intertrial interval. The rotation speed gradually increased from 5 to 80 RPM over 5 minutes. Mice were scored for their latency to fall and the average/best score across three trials was reported. For the pole test, mice first trained for two consecutive days prior to the actual test day. Mice were placed on the top of the pole (7.5 cm from the top of the pole, a 75-cm long, 9-mm diameter metal rod wrapped in bandage gauze) facing the head-up. Total time taken to reach the base of the pole was recorded. Each session consisted of three test trials. The maximum cutoff time to stop the test and recording was 60 s. One mouse (αsyn: n = 19, CTL: n = 20) reached this cutoff and was excluded from the analysis.

### Cognitive Behavioral Assessment

#### Instrumental learning

A detailed protocol for operant training (DOI: dx.doi.org/10.17504/protocols.io.4r3l22k93l1y/v2) can be found here: https://www.protocols.io/view/basic-operant-behavioral-training-4r3l22k93l1y/v1. For operant training, animals were food deprived to 90% of their ad libitum weight. Operant behavior was conducted in custom-made 18 x 18 x 26 cm operant chambers, enclosed within sound-attenuating cabinets (Med Associates Inc.). One side of each operant chamber was equipped with a yellow LED house-light and remained illuminated during all experimental stages, unless stated otherwise. The opposite side of the chamber was equipped with two side ports (left and right) 12 cm apart. Each side port had two yellow LED stimulus lights: one positioned 6 cm above and another inside the port. At a central port positioned between the side ports, sweetened condensed milk (diluted 1:3 with water) was delivered with a solenoid valve (Lee Company). An infrared detector was mounted horizontally across each port to detect nosepoke entries. Experimental events were managed by a PC running Arduino software. The Arduino scripts for running training and testing phases are available here: https://github.com/UCSF-Nelson-Lab/ICD-Cognitive-Arduino.

Operant training began three weeks after αsyn or CTL AAV injections, using a three-phase shaping protocol. Each mouse was trained in the same operant chamber throughout the study. Male and female mice were trained in separate operant chambers. To promote consistency, testing was performed at the same time of day, five days a week.

In Phase 1 (Magazine training, 1 session), food-restricted mice were placed in the operant box, and liquid reward (milk, 10 μL) was delivered at random 40-80s intervals for a total of 40 rewards. Each trial began with the delivery of milk into the center port, accompanied by a 10-second illumination of the center port light. Animals could retrieve the milk reward at any time.

In Phase 2 (Blocked instrumental learning, 6-12 sessions), mice were trained to nosepoke at the side ports to obtain reward on a fixed-ratio (FR1) schedule. Each training session was separated into blocks, separated by a two-minute break period with the house light turned off. Each block lasted 50 minutes. Only nosepokes at one side port (either left or right) were rewarded in each block, and the active side port switched during the break period. The initial active side port (left or right) was counterbalanced across sessions. Each trial began with a stimulus cue light above and within the rewarded port. If the mouse entered that side port within 20 seconds, the cue light was extinguished, and reward (milk, 10 μL) was delivered to the center port, which was illuminated for 10 seconds, followed by a random 30-50 seconds intertrial interval. If the mouse did not enter the correct side port within 20 seconds, cue lights turned off and a timeout of 30-50 seconds began. Mice were trained in Phase 2 for at least 6 days (sessions) and a maximum of 12 days until they reached criterion (correct response on > 80% of trials). Mice that did not meet criterion after 12 days were removed from the study. Overall, 8 out of 19 αsyn mice were excluded during Phase 2 at 3 weeks and 8 out of 20 αsyn mice at 8 weeks. In CTL, 1 out of 20 CTL mice were excluded during Phase 2 at 3 weeks and 1 out 13 CTL mice at 8 weeks.

Phase 3 (Randomized instrumental learning, 6-12 sessions) resembled Phase 2 with two modifications: the introduction of self-initiated trial start, and pseudorandomized (no more than two consecutive trials with the same side cued), rather than blocked trials. Trials began with the center port being illuminated for 10 seconds. During this period, a center nosepoke initiated a trial. Failure to make a center nosepoke within 10 seconds was considered a center omission and resulted in a timeout of 30-50 seconds. Following trial initiation, a side cue light (either left or right) was activated for up to 10 seconds. If the mouse nosepoked the cued side within 10 seconds, the cue light turned off, and reward (milk, 10 μL) was delivered to the center port, which was illuminated for 10 seconds, followed by a 30-50 second intertrial interval. Failure to nosepoke the correct side port within 10 seconds was considered a side omission, and cue lights turned off until next trial. Entering the opposite, uncued side was recorded as an incorrect response, and triggered reward omission. Each session lasted 90 minutes. Mice were trained in Phase 3 for at least 6 days and a maximum of 12 days until criterion was met (correct response on >80% of trials). Mice that did not meet criterion in 12 days were excluded. Overall, 2 out of 12 αsyn mice and 1 out of 17 CTL mice were excluded during Phase 3.

#### Probabilistic reversal learning

A simplified operant training protocol was used to assess probabilistic reversal learning and the entire detailed protocol (DOI: dx.doi.org/10.17504/protocols.io.q26g7p3n9gwz/v1) can be found here: https://www.protocols.io/view/set-shifting-asap-operant-behavior-lernerlab-q26g7p3n9gwz/v1?step=10. Operant training began eight weeks after αsyn or CTL AAV injections, using a simplified shaping protocol.

For this simpler version of operant training, animals were food deprived overnight prior to beginning magazine training, to 85% of their free feeding weight. Operant behavior was conducted in Med Associates Inc. operant box (Operant Box, Med Associates Inc. MED-307W-D2). JN_ASAP_A_MagTraining_30min was loaded into Med Associates and the program was started. House light, fan, corner lights, and cue lights above the left and right nosepoke ports were turned on. On average, every 60 seconds a noncontingent reward of ∼20μl of sweetened condensed milk (diluted 1:3 with water) was delivered via a Med Associates Single Speed Syringe Pump (Liquid Pump, Med Associates, PHM-100). The reward must have been consumed (or a port entry must have been made) for the next reward to be delivered. Session lasted 30 minutes in total.

In FR1 Both (2-12 sessions), mice were trained to initiate trial and nosepoke at either side ports to obtain reward on a fixed-ratio (FR1) schedule. After JN_ASAP_B_FR1_BOTH program was loaded, fan and near-infrared lights were turned on. Each trial had to be initiated by the animal’s nosepoke in the reward port for 140-300 ms. Once the trial was initiated, both left and right cue lights were illuminated and nosepoke on either side led to reward delivery. Reward delivery (milk, 20μl) was associated to a “correct” tone for 0.5s. Each trial had a 3 second intertrial interval and each session lasted 90 minutes or 100 rewards. Mice were trained in FR1 Both for a minimum of 2 days (sessions) and until they reached criterion (correct response on > 80% of trials over 2 consecutive days). Overall, no mice were excluded from the study (αsyn: n= 12, CTL: n=9).

In FR1 Right/FR1 Left (1 session each), mice were trained to follow a light cue and nosepoke at the specific corresponding side port to obtain reward. After either JN_ASAP_C_FR1_RIGHT or JN_ASAP_C_FR1_LEFT programs were loaded, fan and near-infrared lights were turned on. Each session started with the side on which the animal earned more rewards during the previous phase (FR1 Both). Each trial had to be initiated by the animal’s nosepoke in the reward port for 140-300 ms. Once the trial was initiated, either left or right cue lights were illuminated (depending on which program was loaded) and only nosepoke on the illuminated side port led to reward delivery. Reward delivery (milk, 20μl) was associated to “correct” tone for 0.5s. If nosepoke was made on the wrong side port (‘incorrect’ side), the house light illuminated for 7 seconds (‘incorrect’ response) and no reward was delivered. Each trial had a 3 second intertrial interval and each session lasted 90 minutes or 100 rewards. Sessions were conducted daily, alternating between the right and left side regardless of the animal’s performance.

Probabilistic Reversal Learning contained two phases. In the initial phase (deterministic 100/0; 6-14 sessions), the two side ports were associated with 100 (high) and 0% (low) probability of reward, and contingencies were switched every 40 rewarded trials. In the second phase (probabilistic: 90/10; 6-12 sessions), uncertainty increased and the two choice ports were associated with 90 (high) and 10% (low) probability of reward, and contingencies were switched every 40 rewarded trials. Each session lasted 90 minutes with no maximum number of rewards or block switches. The left and right side ports were randomly assigned to either “high” or “low” probability outcomes.

After JN_ASAP_D_100_0_40RewardsSwitch program was loaded, fan and near-infrared lights were turned on. Each trial had to be initiated by the animal’s nosepoke in the reward port for 140-300 ms. Once the trial was initiated, both cue lights were illuminated. Left or right was randomly selected to begin as the high probability side. Only nosepokes on the high probability (100%) side led to a reward. When a nosepoke was made on the high probability side, a ∼20μl of the sweetened condensed milk rewards was delivered associated to a 0.5 second ‘correct’ tone. When a nosepoke was made on the low probability side (0%), the house light illuminated for 7 seconds to signify an incorrect response and no reward was delivered. After reward delivery, a 3 second timeout period started. If the animal earned 40 rewards, a new block began where the high probability (100%) and low probability (0%) sides were switched. If the animal earned 40 rewards on the new high probability side, a new block began where the high and low probability sides switched once again. Each session lasted 90 minutes. There was no maximum number of delivered rewards or block switches. After 3 consecutive days of >80% accuracy (Left Rewards+Right Rewards/Total Trials x 100) or 14 days, animals moved on to the next program.

JN_ASAP_F_90_10_50RewardsSwitch program (2-10 sessions) had a similar structure as the one described above. Once the trial was initiated, both cue lights were illuminated and left or right was randomly selected to begin as the high probability side. When a nosepoke was made on the high probability side, reward (milk, 20μl) and 0.5 second ‘correct’ tone were delivered 90% of the time when the house light illuminated for 7 seconds and no reward was delivered 10 % of the time. If the animals completed 50 trials on the high probability side (90%), a new block started where the high probability (90%) and low probability (10%) sides were switched. If the animal completed 50 trials on the new high probability side, a new block began where the high and low probability sides switch once again until session ended (90 minutes). Mice reached criteria with >70% accuracy (Left Rewards + Right Rewards / Total trials x 100) or a maximum of 10 days. Overall, no mice were excluded from the study (αsyn: n= 12, CTL: n=9).

### Histology

Detailed protocols for preparation of histological sections (DOI: https://www.protocols.io/view/immunohistochemistry-14egn7nezv5d/v1) and semi-quantitative scoring (DOI: https://dx.doi.org/10.17504/protocols.io.81wgbrmknlpk/v1) of midbrain mCherry histology can be found here https://www.protocols.io/view/immunohistochemistry-14egn7nezv5d/v1 and https://www.protocols.io/view/semi-quantitative-histology-scoring-81wgbrmknlpk/v1. After behavioral experiments were completed, mice were deeply anesthetized with I.P. ketamine-xylazine and transcardially perfused with 4% paraformaldehyde in PBS. After perfusion, the brain was dissected from the skull and post-fixed overnight in 4% paraformaldehyde, then placed in 30% sucrose with 0.01% sodium azide at 4°C. The brain was then cut into 30 μm coronal sections on a freezing microtome (Leica) and placed in a 1X PBS-containing well-plate for immunostaining.

Triple immunostaining was performed to assess the expression of (1) human αsyn, (2) mCherry, and (3) tyrosine hydroxylase (TH) expression in the midbrain and striatum. Tissue was blocked with 3% normal donkey serum (NDS) and permeabilized with 0.1% Triton X-100 for 2 hours at room temperature on a shaker. Sections were then incubated in 1X PBS containing primary antibodies and 3% NDS for two nights at 4°C on a shaker.

Primary antibodies used: rabbit anti-mCherry (Abcam, Lot No. ab167453, 1:500), chicken anti-TH (Peel-Freez biologicals, Lot No. P40101-150, 1:1000), rat anti-alpha-synuclein (Enzo Life Sciences, Lot No. ALX-804-258L001, 1:300). Slices were washed 3x with 1X PBS and then incubated in 3% NDS with secondary antibodies for 2 overnights at 4 ° C on a shaker. Secondary antibodies used: Donkey or Goat anti-rabbit Alexa 568 (Jackson ImmunoResearch Laboratories, Lot No. 168192, 1:300), donkey anti-rat Alexa 488 (Jackson ImmunoResearch Laboratories, Lot No. 173054, 1:300) and donkey anti-chicken Alexa 647 (Jackson ImmunoResearch Laboratories, Lot No. 165153, 1:500). Sections were washed 3-5 times with 1X PBS and mounted onto slides for imaging.

Images at 20x magnification for SNc and striatal sections were acquired using a slide scanner microscope (Axioscan 7, Zeiss). Consistent lighting and exposure settings were maintained for all slides for reliable comparison and subsequent image analysis. The Mouse Brain Atlas in Stereotaxic Coordinates (hard copy, 3^rd^ edition) was consulted for anatomical reference.

To assess αsyn and mCherry expression in midbrain dopaminergic neurons, mCherry staining was scored manually by two blinded raters on a scale from 1 to 5. The proportion of neurons stained for mCherry was compared to the total number of midbrain dopaminergic neurons, using TH immunostaining. A score of 1 represented 0-20%; 2: 20-40%; 3: 40-60%; 4: 60-80% and 5: 80-100%. The score was based on mCherry expression, so it could be used in both CTL and αsyn mice. Synuclein expression was also confirmed in αsyn mice in a later step. Zeiss Microscopy software or QuPath-0.7.0 software was used for visualization. For each mouse, 4-6 coronal sections taken from -3.88mm to -2.54mm AP were analyzed and each given an individual score. The average score was computed from all coronal sections. Mice were included if the histological score was 2 or greater on both sides of the midbrain. If scores between sides differed by >2, the animal was excluded. Overall, 20 out of 68 αsyn mice and 20 out of 65 CTL mice were excluded based on mCherry expression.

## EXPERIMENTAL DESIGN AND STATISTICAL ANALYSIS

The experimental design and statistical analysis of all key experiments are summarized in Table 1, Supplemental Information. This table includes statistical tests, *N* (animals), *p* values and the associated figures. All data are presented as the mean ± SEM. The statistical tests were performed using Python or R.

## AUTHOR CONTRIBUTIONS

BC, XZ, JS, JN, TL and ABN designed experiments. BC, XZ, JS, JN, KR performed experiments and analyses. BC and ABN wrote the manuscript, with input from all authors. ABN and TL obtained funding.

## ACKOWLEDGMENTS.

This work was supported by a grant from Aligning Science Across Parkinson’s (ASAP020529 to A.B.N., R. E., G.H., and T.L.) through the Michael J. Fox Foundation for Parkinson’s Research (MJFF). For the purpose of open access, the author has applied a CC BY public copyright license to all Author Accepted Manuscripts arising from this submission. We are grateful to all the members of the Nelson and Lerner Labs, as well as Chris Ford, Kelsey Barcomb, Vijay Namboodiri, Zayd Khaliq, and Ken Nakamura for valuable suggestions and feedback on this manuscript.

## DATA AVAILABILTY

Datasets generated during the present study have been deposited on Zenodo and are publicly available (10.5281/zenodo.24081901). Detailed protocols and are listed within the Methods section.

## CODE AVAILABILITY

The original codes generated for the present study are available in Zenodo and are publicly available as of the date of publication (https://doi.org/10.5281/zenodo.24365624).

