## Supplemental Figures and Tables for "Overexpression of α−synuclein in Midbrain Dopamine Neurons Drives Learning Deficits in Mice"

SUPPLEMENTARY INFORMATION

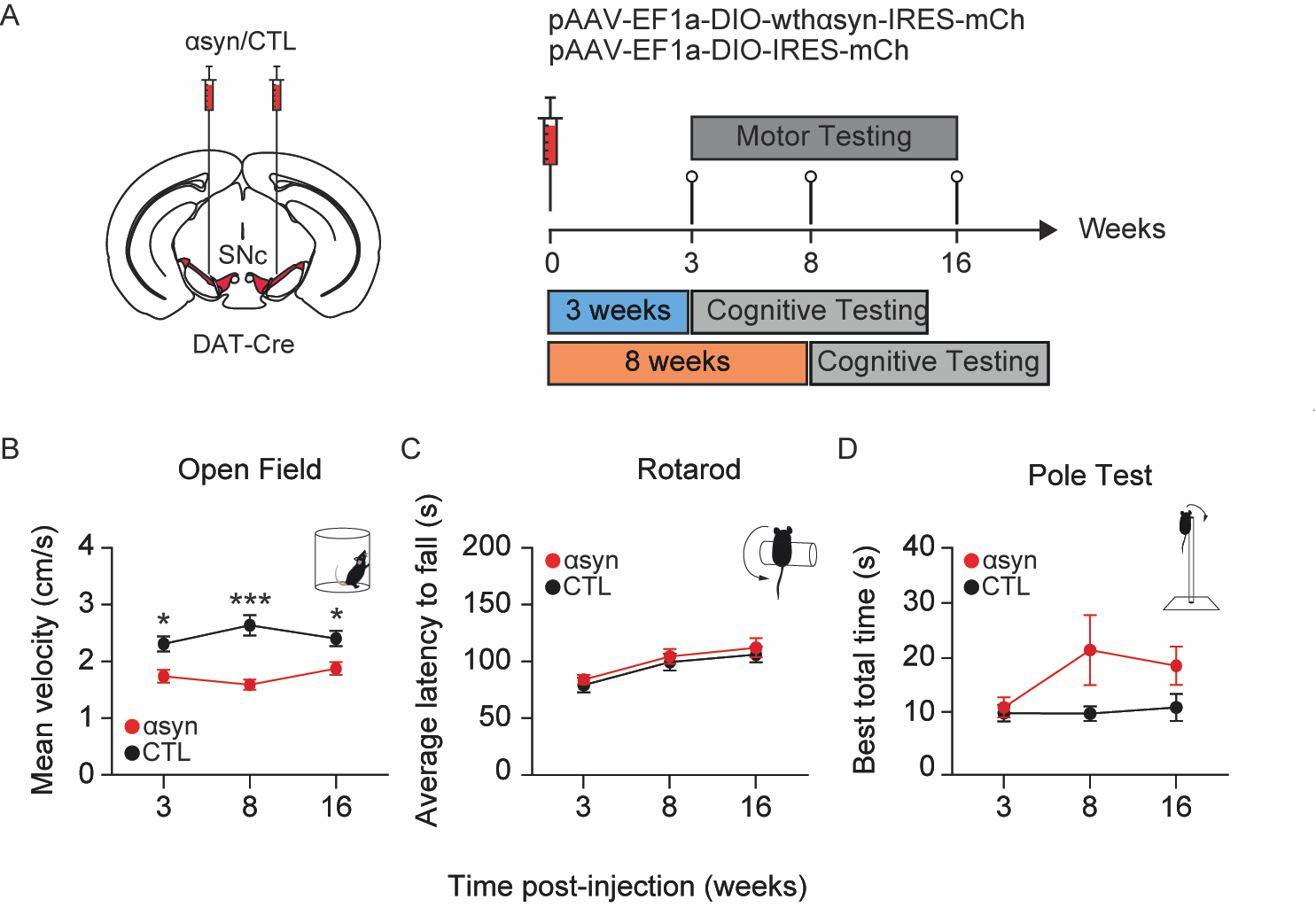

**Supplementary Figure 1: α-synuclein overexpression in midbrain dopamine neurons induces subtle and stable motor deficits at 3, 8, and 16 weeks.**

**A.** Left: AAVs encoding human wild-type alpha synuclein (αsyn) and/or mCherry (CTL) were injected into the midbrain of DAT-Cre mice. SNc: Substantia nigra pars compacta.. Right: Experimental timeline. **B.** Mean velocity (cm/s) in αsyn and CTL mice, at 3 (n=24, 26), 8 (n=28, 28), and 16 (n=21, 23) weeks post-injection during open field test. A linear mixed-effects model showed a significant effect over time at 3 (Day: p = 0.0082), 8 (Day: p = < 0.0001), and 16 (Day: p = 0.0221) weeks. Data are presented as mean ± SEM. See Supplementary Information, Table 1 for statistics. **C.** Latency to fall (s) in αsyn and CTL mice, at 3 (n=24, 26), 8 (n=21, 25), and 16 (n=17, 21) weeks post-injection during accelerating rotarod test. A linear mixed-effects model showed a significant effect over time at 3 (Day: p = 0.5961), 8 (Day: p = 0.8338), and 16 (Day: p = 0.5473) weeks. Data are presented as mean ± SEM. See Supplementary Information, Table 1 for statistics. **D.** Best total time (s) in αsyn and CTL mice, at 3 (n=10, 9), 8 (n=6, 6), and 16 (n=5, 6) weeks post-injection. A linear mixed-effects model showed a significant effect over time at 3 (Day: p = 0.1709), 8 (Day: p = 0.6174), and 16 (Day: p = 0.7683) weeks. Data are presented as mean ± SEM. See Supplementary Information, Table 1 for statistics.

Table 1. EXPERIMENTAL ANALYSIS AND STATISTICS

| **Table 1. EXPERIMENTAL ANALYSIS AND STATISTICS**  Linear mixed-effects models (LMMs) with Type III ANOVA; two-way repeated-measures ANOVA; Welch’s two-sample t-tests, unpaired two-tailed Student’s t-test | | | | | | | | | |
| --- | --- | --- | --- | --- | --- | --- | --- | --- | --- |
| **Key Experiments** | **Figure** | **Comparison** | **Test** | **Statistic** | **df** | **N** (CTL) | **N** (αsyn) | ***p* value** |  |
| Learning Curve, 3 weeks | Figure 1E, left panel | Condition | LMM, Type III ANOVA | F = 1.9712 | 36.999 | 20 | 19 | 0.1686 | ns |
|  |  | Day | LMM, Type III ANOVA | F = 45.0335 | 182.053 | 20 | 19 | <0.0001 | *** |
|  |  | Condition x Day | LMM, Type III ANOVA | F = 2.4751 | 182.053 | 20 | 19 | 0.0338 | * |
| Learning Curve, 3 weeks | Figure 1E, left panel | Day 1: αsyn vs CTL | Pairwise, Holm-adjusted | t = -0.491 | 56.9 | 20 | 19 | 0.6250 | ns |
|  |  | Day 2: αsyn vs CTL | Pairwise, Holm-adjusted | t = -0.651 | 56.4 | 20 | 19 | 0.5177 | ns |
|  |  | Day 3: αsyn vs CTL | Pairwise, Holm-adjusted | t = -0.779 | 56.9 | 20 | 19 | 0.4391 | ns |
|  |  | Day 4: αsyn vs CTL | Pairwise, Holm-adjusted | t = -1.298 | 56.9 | 20 | 19 | 0.1995 | ns |
|  |  | Day 5: αsyn vs CTL | Pairwise, Holm-adjusted | t = -1.844 | 56.9 | 20 | 19 | 0.0704 | ns |
|  |  | Day 6: αsyn vs CTL | Pairwise, Holm-adjusted | t = -2.470 | 56.9 | 20 | 19 | 0.0165 | * |
| Learning Curve, 8 weeks | Figure 1E, right panel | Condition | LMM, Type III ANOVA | F = 23.837 | 31.014 | 13 | 20 | <0.0001 | *** |
|  |  | Day | LMM, Type III ANOVA | F = 32.849 | 154.047 | 13 | 20 | <0.0001 | *** |
|  |  | Condition x Day | LMM, Type III ANOVA | F = 10.581 | 154.047 | 13 | 20 | <0.0001 | *** |
| Learning Curve, 8 weeks | Figure 1E, right panel | Day 1: αsyn vs CTL | Pairwise, Holm-adjusted | t = -1.244 | 61.3 | 13 | 20 | 0.2182 | ns |
|  |  | Day 2: αsyn vs CTL | Pairwise, Holm-adjusted | t = -2.405 | 61.3 | 13 | 20 | 0.0192 | * |
|  |  | Day 3: αsyn vs CTL | Pairwise, Holm-adjusted | t = -3.938 | 61.3 | 13 | 20 | 0.0002 | *** |
|  |  | Day 4: αsyn vs CTL | Pairwise, Holm-adjusted | t = -4.868 | 61.3 | 13 | 20 | <0.0001 | *** |
|  |  | Day 5: αsyn vs CTL | Pairwise, Holm-adjusted | t = -5.586 | 61.3 | 13 | 20 | <0.0001 | *** |
|  |  | Day 6: αsyn vs CTL | Pairwise, Holm-adjusted | t = -6.409 | 61.3 | 13 | 20 | <0.0001 | *** |
| Reaction Time, 3 weeks | Figure 2A, left panel | Condition | LMM, Type III ANOVA | F = 0.0335 | 35.81 | 20 | 19 | 0.8558 | ns |
|  |  | Day | LMM, Type III ANOVA | F = 5.8030 | 160.29 | 20 | 19 | <0.0001 | *** |
|  |  | Condition x Day | LMM, Type III ANOVA | F = 0.8326 | 160.29 | 20 | 19 | 0.5283 | ns |
| Reaction Time, 8 weeks | Figure 2A, right panel | Condition | LMM, Type III ANOVA | F = 0.2843 | 27.879 | 13 | 20 | 0.5981 | ns |
|  |  | Day | LMM, Type III ANOVA | F = 4.0755 | 121.388 | 13 | 20 | <0.01 | * |
|  |  | Condition x Day | LMM, Type III ANOVA | F = 0.7507 | 121.388 | 13 | 20 | 0.5871 | ns |
| Central nosepoke number, 3 weeks | Figure 2B, left panel | Condition | LMM, Type III ANOVA | F = 0.5175 | 32.071 | 20 | 19 | 0.4771 | ns |
|  |  | Day | LMM, Type III ANOVA | F = 3.4226 | 156.181 | 20 | 19 | < 0.01 | * |
|  |  | Condition x Day | LMM, Type III ANOVA | F = 1.4669 | 156.181 | 20 | 19 | 0.2037 | ns |
| Central nosepoke number, 8 weeks | Figure 2B, right panel | Condition | LMM, Type III ANOVA | F = 9.1028 | 24.026 | 13 | 20 | <0.01 | ** |
|  |  | Day | LMM, Type III ANOVA | F = 3.0347 | 119.097 | 13 | 20 | 0.0129 | * |
|  |  | Condition x Day | LMM, Type III ANOVA | F = 0.3832 | 119.097 | 13 | 20 | 0.8595 | ns |
| % side omissions, 3 weeks | Figure 2C, left panel | Condition | LMM, Type III ANOVA | F = 1.9109 | 36.993 | 20 | 19 | 0.1751 | ns |
|  |  | Day | LMM, Type III ANOVA | F = 44.6781 | 182.047 | 20 | 19 | <0.0001 | *** |
|  |  | Condition x Day | LMM, Type III ANOVA | F = 2.3988 | 182.047 | 20 | 19 | 0.0390 | * |
| % side omissions, 3 weeks | Figure 2C, left panel | Day 1: αsyn vs CTL | Pairwise, Holm-adjusted | t = 0.477 | 56.9 | 20 | 19 | 0.6353 | ns |
|  |  | Day 2: αsyn vs CTL | Pairwise, Holm-adjusted | t = 0.660 | 59.4 | 20 | 19 | 0.5117 | ns |
|  |  | Day 3: αsyn vs CTL | Pairwise, Holm-adjusted | t = 0.798 | 56.9 | 20 | 19 | 0.4284 | ns |
|  |  | Day 4: αsyn vs CTL | Pairwise, Holm-adjusted | t = 1.210 | 56.9 | 20 | 19 | 0.2313 | ns |
|  |  | Day 5: αsyn vs CTL | Pairwise, Holm-adjusted | t = 1.837 | 56.9 | 20 | 19 | 0.0714 | ns |
|  |  | Day 6: αsyn vs CTL | Pairwise, Holm-adjusted | t = 2.436 | 56.9 | 20 | 19 | 0.0180 | * |
| % side omissions, 8 weeks | Figure 2C, right panel | Condition | LMM, Type III ANOVA | F = 23.515 | 31.014 | 13 | 20 | <0.0001 | *** |
|  |  | Day | LMM, Type III ANOVA | F = 32.043 | 154.048 | 13 | 20 | <0.0001 | *** |
|  |  | Condition x Day | LMM, Type III ANOVA | F = 10.111 | 154.048 | 13 | 20 | <0.0001 | *** |
| % side omissions, 8 weeks | Figure 2C, right panel | Day 1: αsyn vs CTL | Pairwise, Holm-adjusted | t = 1.281 | 61.9 | 13 | 20 | 0.2049 | ns |
|  |  | Day 2: αsyn vs CTL | Pairwise, Holm-adjusted | t = 2.366 | 61.9 | 13 | 20 | 0.0211 | * |
|  |  | Day 3: αsyn vs CTL | Pairwise, Holm-adjusted | t = 3.883 | 61.9 | 13 | 20 | 0.0003 | *** |
|  |  | Day 4: αsyn vs CTL | Pairwise, Holm-adjusted | t = 4.901 | 62.8 | 13 | 20 | <0.0001 | *** |
|  |  | Day 5: αsyn vs CTL | Pairwise, Holm-adjusted | t = 5.403 | 61.9 | 13 | 20 | <0.0001 | *** |
|  |  | Day 6: αsyn vs CTL | Pairwise, Holm-adjusted | t = 6.386 | 61.9 | 13 | 20 | <0.0001 | *** |
| Average latency (s), 3 weeks | Figure 2D, left panel | CTL vs αsyn | Welch’s *t*-test | t = 0.81676 | 38.569 | 20 | 22 | 0.4191 | ns |
| Average latency (s), 8 weeks | Figure 2D, right panel | CTL vs αsyn | Welch’s *t*-test | t = 0.59448 | 17.503 | 12 | 16 | 0.5598 | ns |
| Motivation Test | Figure 2E | CTL vs αsyn | Unpaired two-tailed Student’s *t*-test | t = -0.097656 | 24 | 11 | 15 | 0.923 | ns |
| Learning Curve, 3 weeks | Figure 3B | Condition | LMM, Type III ANOVA | F = 0.1549 | 26.93 | 17 | 12 | 0.6970 | ns |
|  |  | Day | LMM, Type III ANOVA | F = 4.5600 | 133.96 | 17 | 12 | <0.0001 | *** |
|  |  | Condition x Day | LMM, Type III ANOVA | F = 0.6865 | 133.96 | 17 | 12 | 0.6344 | ns |
| Days to criterion | Figure 4C, 4E | Training Stage | Two-way RM ANOVA, GG corrected | F = 32.586 | 35.215 | 12 | 9 | <0.0001 | *** |
|  |  | Condition | Two-way RM ANOVA, GG corrected | F = 13.930 | 19 | 12 | 9 | 0.0014 | ** |
|  |  | Condition x Training Stage | Two-way RM ANOVA, GG corrected | F = 1.348 | 38 | 12 | 9 | 0.2720 | ns |
| Days to criterion | Figure 4C | FR1 | Bonferroni–Šídák multiple comparison | F = 3.173 | 10.520 | 12 | 9 | 0.0278 | * |
|  | Figure 4E | 100/0 | Bonferroni–Šídák multiple comparison | F = 2.314 | 18.371 | 12 | 9 | 0.0943 | ns |
|  | Figure 4E | 90/10 | Bonferroni–Šídák multiple comparison | F = 0.864 | 12.686 | 12 | 9 | 0.7879 | ns |
| % correct after switch, 100/0 | Figure 4G, left panel | Trials | Two-way RM ANOVA, GG corrected | F = 80.56 | 109.68 | 12 | 9 | <0.0001 | *** |
|  |  | Condition | Two-way RM ANOVA | F = 3.782 | 19 | 12 | 9 | 0.0668 | ns |
|  |  | Trials x Condition | Two-way RM ANOVA | F = 2.762 | 456 | 12 | 9 | <0.0001 | *** |
| % correct after switch, 90/0 | Figure 4G, right panel | Trials | Two-way RM ANOVA, GG corrected | F = 81.63 | 122.38 | 12 | 9 | <0.0001 | *** |
|  |  | Condition | Two-way RM ANOVA | F = 4.777 | 19 | 12 | 9 | 0.0416 | * |
|  |  | Trials x Condition | Two-way RM ANOVA | F = 0.989 | 456 | 12 | 9 | 0.4791 | ns |
| Velocity (cm/sec) | Supplementary Figure 1B | Condition | LMM, Type III ANOVA | F = 19.7883 | 45.013 | 31 | 29 | <0.0001 | *** |
|  |  | Age | LMM, Type III ANOVA | F = 1.5458 | 80.936 | 31 | 29 | 0.2193 | ns |
|  |  | Condition x Age | LMM, Type III ANOVA | F = 5.1637 | 80.936 | 31 | 29 | 0.0078 | * |
| Velocity (cm/sec) | Supplementary Figure 1B | 3 weeks: αsyn vs CTL | Pairwise, Holm-adjusted | t = -2.689 | 114 | 24 | 26 | 0.0082 | * |
|  |  | 8 weeks: αsyn vs CTL | Pairwise, Holm-adjusted | t = -5.726 | 107 | 28 | 28 | <0.0001 | *** |
|  |  | 16 weeks: αsyn vs CTL | Pairwise, Holm-adjusted | t = -2.318 | 122 | 21 | 23 | 0.0221 | * |
| Latency to fall (s) | Supplementary Figure 1C | Condition | LMM, Type III ANOVA | F = 0.3623 | 42.862 | 24 | 26 | 0.5504 | ns |
|  |  | Age | LMM, Type III ANOVA | F = 15.9553 | 79.030 | 24 | 26 | <0.0001 | *** |
|  |  | Condition x Age | LMM, Type III ANOVA | F = 0.0737 | 79.030 | 24 | 26 | 0.9290 | ns |
| Latency to fall (s) | Supplementary Figure 1C | 3 weeks: αsyn vs CTL | Pairwise, Holm-adjusted | t = 0.532 | 102 | 24 | 26 | 0.5961 | ns |
|  |  | 8 weeks: αsyn vs CTL | Pairwise, Holm-adjusted | t = 0.8338 | 107 | 21 | 25 | 0.8338 | ns |
|  |  | 16 weeks: αsyn vs CTL | Pairwise, Holm-adjusted | t = 0.5473 | 117 | 17 | 21 | 0.5473 | ns |
| Total Time (s) | Supplementary Figure 1D | Condition | LMM, Type III ANOVA | F = 1.1240 | 1.1240 | 10 | 10 | 0.3087 | ns |
|  |  | Age | LMM, Type III ANOVA | F = 0.4989 | 22.428 | 10 | 10 | 0.6138 | ns |
|  |  | Condition x Age | LMM, Type III ANOVA | F = 0.2530 | 22.428 | 10 | 10 | 0.7786 | ns |
| Total Time (s) | Supplementary Figure 1D | 3 weeks: αsyn vs CTL | Pairwise, Holm-adjusted | t = 1.396 | 37.7 | 10 | 9 | 0.1709 | ns |
|  |  | 8 weeks: αsyn vs CTL | Pairwise, Holm-adjusted | t = 0.504 | 38.8 | 6 | 6 | 0.6174 | ns |
|  |  | 16 weeks: αsyn vs CTL | Pairwise, Holm-adjusted | t = 0.297 | 39.0 | 5 | 6 | 0.7683 | ns |

Table 2. KEY RESOURCE TABLE

| **REAGENT or RESOURCE** | **SOURCE** | **IDENTIFIER** |
| --- | --- | --- |
| **Antibodies** | | |
| Rabbit Anti-mCherry | Abcam | Cat# ab167453, RRID: AB_2571870 |
| Chicken Anti-  Tyrosine Hydroxylase (TH) | Peel-Freez biologicals | Cat# P40101-150, RRID: AB_2617184 |
| Rat Anti-Alpha  Synuclein | Enzo Life  Sciences | Cat# ALX-804-258L001, RRID: AB_2050691 |
| Alexa Fluor 568-Goat Anti-Rabbit IgG | Jackson ImmunoResearch | Cat# 111-575-144, RRID: AB_3095432 |
| Alexa Fluor 568-Donkey Anti-Rabbit IgG | Jackson ImmunoResearch | Cat#, 711-575-152, RRID: AB_3095472 |
| Alexa Fluor 647-Donkey Anti-Chicken IgG | Jackson ImmunoResearch | Cat# 703-605-155, RRID: AB_2340379, Lot #165153 |
| **Bacterial and Virus Strains** | | |
| AAV5-Ef1a-DIO-wthasyn-IRES-mCh | Vector Builder | Lot # 211030AAVJ01, RRID: 264940 |
| AAV5-Ef1a-DIO-IRES-mCh | Vector Builder | Lot # 211104AAVD11, RRID: 264939 |
| **Chemicals, Peptides, and Recombinant Proteins** | | |
| VECTASHIELD Antifade Mounting Medium | Vector Laboratories | RRID: AB_2336789; Cat# H-1000 |
| **Experimental Models: Organisms/Strains** |  |  |
| Mouse: B6.SJL-Slc6a3^tm1.1(cre)Bkmn^/J | The Jackson Laboratory | Strain:#006660, RRID:IMSR_JAX:006660 |
| Mouse: C57Bl/6J | The Jackson Laboratory | Strain:#000664,  RRID: IMSR_JAX:000664 |
| **Software and Algorithms** |  |  |
| EthoVision XT | Noldus | <https://www.noldus.com/animal-behavior-research/products/ethovision-xt>; RRID: SCR_000441 |
| Adobe Illustrator CS5 | Adobe | <https://www.adobe.com/products/illustrator.html>; RRID: SCR_014198 |
| QuPath-0.7.0 | QuPath | <https://qupath.github.io/>; RRID:SCR_018257 |
| Spyder | Spyder | <https://www.spyder-ide.org/>;  RRID:SCR_017585 |
| Anaconda | Anaconda | <https://docs.anaconda.com/> ;  RRID:SCR_025572 |
| GraphPad Prism Version 10 | GraphPad | <https://www.graphpad.com/>; RRID:SCR_002798 |
| [MED-PC](https://rrid.site/data/record/nlx_144509-1/SCR_012156/resolver?q=%22MED%20PC%22&l=%22MED%20PC%22&i=rrid:scr_012156) 5 | Med Associates, Inc. | <https://med-associates.com/product/med-pc/>; RRID:SCR_012156 |
| **Other** |  |  |
| Mouse operant chamber | Med Associates, Inc. | <https://med-associates.com/>;  MED-307W-D2 |
| Syringe pump | Med Associates, Inc. | <https://med-associates.com/>;  PHM-100 |
